# Transcriptomic profiling of normal and glaucomatous trabecular meshwork cells in a tissue-like soft ECM environment

**DOI:** 10.64898/2026.10.26.761819

**Authors:** Rajanya Ghosh, Rachel Hadvina, Ana N. Strat, Preethi S. Ganapathy, W. Daniel Stamer, Kate E. Keller, Yutao Liu, Samuel Herberg

## Abstract

**Purpose:** Trabecular meshwork (TM) dysfunction contributes to the elevated intraocular pressure in primary open-angle glaucoma. To better understand the “molecular signature” distinguishing glaucomatous from normal TM cells, we interrogated their genome-wide transcriptomic differences in a tissue-like soft extracellular matrix (ECM) microenvironment.

**Methods:** Primary human glaucomatous and normal TM cell strains (N = 5 each) from three independent laboratories were plated on bioengineered collagen type I/hyaluronic acid/elastin-like polypeptide ECM hydrogels in 10% FBS media for 4-5 days, then serum-free media for 2 days. Cell-laden hydrogels were homogenized in TRIzol for RNA isolation, quality assessment, and library preparation, followed by paired-end 100-bp Illumina sequencing. Differential expression was assessed using DESeq2, including glaucoma status, originating laboratory, and donor age/sex in the model. Differentially expressed genes (DEGs) were defined as p < 0.05 and |fold change| ≥ 2, followed by Gene Ontology (GO) enrichment and QIAGEN Ingenuity Pathway Analysis (IPA).

**Results:** In total, 588 DEGs distinguished the glaucomatous from the normal TM transcriptome (393 up-regulated in glaucomatous TM; 195 up-regulated in normal TM, i.e., down-regulated in glaucomatous TM). GO and IPA causal analyses revealed three coordinated disease programs: (i) TGF-β/Wnt-driven ECM remodeling and fibrosis, (ii) NF-κB-associated inflammation, and (iii) a hypoxia/metabolic-stress program linked to p300/CREBBP chromatin co-activation.

**Conclusions:** These data define a transcriptomic signature of glaucomatous TM cells when cultured in a soft physiologic ECM microenvironment, independent of the originating laboratory, with three mechanistic axes (ECM/fibrosis, inflammation, hypoxia/metabolism), and nominate p300 (encoded by *EP300*) as a candidate upstream node integrating these programs.

## Introduction

Primary open-angle glaucoma (POAG) is a leading cause of irreversible vision loss worldwide^1–3^. It is a chronic optic neuropathy due to elevation of intraocular pressure because of increased aqueous humor outflow resistance^2–6^. This outflow resistance arises from pathological cellular and extracellular matrix (ECM) remodeling of the trabecular meshwork (TM)^7–13^. Dysfunction of tissue-resident TM cells is a central driver of disease progression. Strong evidence suggests that the glaucomatous TM exhibits chronic inflammation^14,15^, oxidative stress^16–19^, and fibrotic remodeling^20–22^, resulting in excessive ECM deposition and tissue stiffening^23–27^. However, the genome-wide transcriptional programs underlying the pathological “molecular signature” in glaucomatous TM cells compared to non-glaucomatous TM cells under physiologically relevant culture conditions are insufficiently characterized.

Early microarray studies identified glaucoma-associated gene expression changes in human TM tissue, including altered expression of genes involved in ECM organization, cell adhesion, inflammation, and antioxidant defenses^28,29^. In a direct comparison of healthy and glaucomatous human tissue, a single-nucleus RNA sequencing study reported cell-type-specific changes in ECM and mechanosensitive pathways, inflammatory signaling, and autophagy and mitophagy^30^. These findings suggest that fibrotic remodeling is accompanied by altered inflammatory and stress-response programs in glaucoma. A broad overview of published TM transcriptomic studies is provided in **Supplementary Table 1**.

Direct comparisons of non-glaucomatous, normal TM (NTM) cells and glaucomatous TM (GTM) cells have identified disease-associated transcriptional changes. Bulk RNA-sequencing and proteomic analyses of the SV40-transformed iHTM and GTM3 cell lines implicated ECM-receptor interactions and cell-survival signaling^31^. In primary donor cell strains, Yang et al. used the NanoString nCounter human fibrosis panel to examine 750 genes in NTM and GTM donor cell strains, identifying differences in fibrosis-related gene expression^32^. While this targeted panel did not provide a genome-wide comparison, these findings indicated dysregulated matrix composition and turnover rather than a uniform increase in ECM gene expression^32^. A recent multiomic study of 18 NTM cell strains from seven laboratories (including ours) provides a reference for normal cultured cells without a GTM comparison^33^. Using dexamethasone treatment as a glaucoma-relevant stimulus, transcriptional and chromatin differences have also been characterized^34^. However, such responses may not fully reproduce the disease-associated programs retained by cells from glaucomatous donors.

Tian et al. performed single-cell RNA sequencing of primary human TM cells from non-glaucomatous and glaucomatous donors. Cultured cells showed reduced cellular heterogeneity and differences in gene expression compared with native TM tissue, although NTM and GTM cultures did not differ significantly in the cell-type markers or clusters examined^35^. These findings emphasize the importance of considering the culture environment when investigating disease-associated transcriptional changes. Conventional tissue culture substrates such as plastic and glass are orders of magnitude stiffer (elastic modulus, >1 GPa) than normal human TM tissue, for which the reported mean elastic modulus ranges from 0.9 to 16 kPa^25,36,37^. Atomic force microscopy studies have identified a higher mean elastic modulus in glaucomatous human TM tissue, ranging from 2.0 to 80 kPa^25,36,37^. Thus, it is well-established that substrate stiffness regulates ECM-associated gene expression and mechanotransduction, and the culture substrate alters TM cell contractile responses to transforming growth factor (TGF)-β2^27,38,39^. RNA-sequencing of normal TM cells cultured on dexamethasone-induced cell-derived matrices has also demonstrated that an altered ECM can change the TM transcriptome ^40^. These studies did not directly compare primary NTM and GTM donor cells on a soft ECM substrate. Hence, it remains unclear which matrix-remodeling, inflammatory, and hypoxia/metabolic-stress signatures distinguish donor-derived NTM and GTM cells under these conditions. To this end, a common soft ECM environment can provide a complementary approach to determine which disease-associated transcriptional features still remain detectable after cells are removed from their native tissue environment.

To address this gap, we performed bulk RNA-sequencing to profile primary human NTM and GTM cells from five independent donors per group, obtained from three independent laboratories. All ten cell strains were studied under the same culture conditions on our established biomimetic soft ECM hydrogels with an elastic modulus of 0.42 ± 0.12 kPa, close to the lower range of native TM tissue (0.9-16 kPa^25,36,37^) while simulating a more favorable tissue-mimetic ECM microenvironment compared to supraphysiologically stiff tissue culture plates. Glaucoma status, laboratory of origin, donor age, and donor sex were included in the differential-expression model. Gene ontology and pathway analyses were used to identify affected biological processes and candidate upstream regulators. This study extends prior NTM and GTM transcriptomic comparisons by examining five independent primary donor cell strains per group from three laboratories on a common soft ECM substrate, allowing us to identify glaucoma-associated transcriptional programs retained outside the native diseased matrix.

## Materials and Methods

### 2.1 TM cell isolation and culture

Experiments using human donor eye tissue were approved by the SUNY Upstate Medical University Institutional Review Board (protocol #1211036) and performed in accordance with the tenets of the Declaration of Helsinki for the use of human tissue. Two primary human NTM cell strains (HTM 36, HTM 28) were isolated from normal (non-glaucomatous) donor corneal rims discarded following corneal transplantation surgeries, as recently described^13,38,41^. Three NTM cell strains (NTM 2021-1110, NTM 2022-0794, NTM 2021-1323) and three GTM cell strains (GTM 2022-0949, GTM 2018-1898, GTM 2018-1672) were obtained from Dr. Keller’s laboratory. The remaining two GTM cell strains (GTM 211, GTM 1445) were obtained from Dr. Stamer’s laboratory. All cells were isolated and cultured according to established protocols^42–44^. Except for HTM 28, all NTM and GTM cell strains have been previously characterized/validated with dexamethasone-induced myocilin expression in more than 50% of cells by immunocytochemistry and immunoblot analyses^13,32,38,41,45–49^, as per accepted standards in the field^43^. Detailed characterization for HTM 28 cell is provided in the **Supplementary Information** and **Supplementary Figure 1**. For RNA-seq, five NTM and five GTM cell strains were used with 3 technical replicates each. All studies were conducted between cell passages 3-6. All NTM and GTM cells were cultured in low glucose Dulbecco’s Modified Eagle’s Medium (DMEM; Gibco, Thermo Fisher Scientific) containing 10% fetal bovine serum (FBS; Atlanta Biologicals, Flowery Branch, GA, USA) and 1% penicillin/streptomycin/glutamine (PSG; Gibco), and maintained at 37°C in a humidified atmosphere with 5% CO_2_. Fresh media was supplied every 2-3 days. Of note, due to the mix of TM cell strain nomenclatures used in the different laboratories, we simplified the sample IDs for this study to avoid confusion. Demographics for all NTM and GTM cells from the different laboratories are listed in **Table 1**.

**Table 1.** Demographics for NTM and GTM cell strains.

| Sample ID | Original ID | PI | Age | Sex | Tissue source | Cause of death | Glaucoma | Relevant clinical background / medications | Characterization |
| --- | --- | --- | --- | --- | --- | --- | --- | --- | --- |
| NTM 1 | HTM 36 | SH | 56 y | Female | Corneal rim after surgery (DSAEK) | Unknown | No | Unknown | Bagué et al. 2022<sup>50</sup> |
| NTM 2 | HTM 28 | SH | 64 y | Male | Corneal rim after surgery (DSAEK) | Unknown | No | Unknown | Supplementary Figure 1 |
| NTM 3 | NTM 2022-0794 | KEK | 80 y | Male | Whole globe | Respiratory failure with hypoxia | No | Unknown | Yang et al. 2025<sup>32</sup> |
| NTM 4 | NTM 2021-1110 | KEK | 77 y | Female | Whole globe | Lung cancer | No | Unknown | Yang et al. 2022<sup>48</sup> |
| NTM 5 | NTM 2021-1323 | KEK | 57 y | Male | Whole globe | Cardiac arrest | No | Unknown | Yang et al. 2022<sup>48</sup> |
| GTM 1 | GTM 2022-0949 | KEK | 61 y | Female | Whole globe | Pneumonia | Yes | POAG | Yang et al. 2025<sup>32</sup> |
| GTM 2 | GTM 2018-1898 | KEK | 68 y | Male | Whole globe | Sepsis | Yes | Glaucoma (recent dx), IOL OU | Sun et al. 2019a<sup>47</sup>; Sun et al. 2019b<sup>49</sup>; |
| GTM 3 | GTM 2018-1672 | KEK | 57 y | Male | Whole globe | Respiratory shock | Yes | Glaucoma (dx 2001; tx eye drops per NOK) | Sun et al. 2019a<sup>47</sup>; Sun et al. 2019b<sup>49</sup>; |
| GTM 4 | GTM 1445 | WDS | 81 y | Female | Whole globe | Unknown | Yes | POAG; brimonidine; dorzolamide-timolol; latanoprost | Li et al. 2021<sup>41</sup> |
| GTM 5 | GTM 211 | WDS | 75 y | Female | Whole globe | Unknown | Yes | Glaucoma; bilateral cataract & retina surgery; latanoprost (Xalatan) | Li et al. 2021<sup>41</sup>; Li et al. 2022<sup>13</sup> |

### 2.2 ECM hydrogel preparation

ECM hydrogel precursors methacrylate-conjugated bovine collagen type I (Advanced BioMatrix, Carlsbad, CA, USA; 3.6 mg/ml final), thiol-conjugated hyaluronic acid (Glycosil®; Advanced BioMatrix; 0.5 mg/ml), 0.1% (w/v) photoinitiator LAP (lithium phenyl-2,4,6-trimethylbenzoylphosphinate; Sigma-Aldrich, St. Louis, MO, USA), and in-house-expressed elastin-like polypeptide (thiol via KCTS flanks^51^; 2.5 mg/ml) were thoroughly mixed. ECM hydrogels were prepared by pipetting 1.5 ml of the precursor solution into each well of a 6-well plate (Corning; Thermo Fisher Scientific). Constructs were crosslinked by exposure to blue light (405-500 nm, 320 mW/cm^2^, 10 s; OmniCure S1500 UV Spot Curing System; Excelitas Technologies, Mississauga, Ontario, Canada) as previously described^52^. ECM hydrogel elastic modulus was determined by oscillatory rheology analysis (**Supplementary Information**).

### 2.3 NTM/GTM cell seeding

ECM hydrogels were allowed to equilibrate for 1-1.5 h in DMEM, before NTM/GTM cells were seeded at 2 × 10⁴ cells/cm² atop the pre-formed hydrogels. For the first 4-5 days, cells were cultured in low glucose DMEM containing 10% FBS and 1% PSG until ∼80-90% confluency, after which the media was changed to serum-free low-glucose DMEM containing 1% PSG for 2 days and maintained at 37°C with 5% CO_2_.

### 2.4 RNA sequencing

NTM/GTM cell-laden ECM hydrogels (N = 5 per group) from 2-3 wells of a 6-well plate (Thermo Fisher Scientific) were collected with a spatula into glass test tubes containing 2 ml TRIzol™ (Invitrogen). Samples were homogenized using a tissue homogenizer for 7-8 min, transferred to a 15-ml conical tube, and handed to the SUNY Upstate Medical University Molecular Analysis Core (or stored at-80°C until transfer) for RNA isolation, quality control, and library preparation. RNA concentration and integrity were assessed using an Agilent 2100 Bioanalyzer (Agilent Technologies, Santa Clara, CA, USA); RNA concentration, 28S/18S rRNA ratio, and RIN for all sequenced samples are reported in **Supplementary Table 2**. Strand-specific RNA-seq libraries were prepared from 100 nanograms of total RNA per sample using the Illumina Stranded Total RNA Prep, Ligation with Ribo-Zero Plus kit (Illumina, San Diego, CA, USA), which removes ribosomal RNA by enzymatic depletion. Paired-end 100-bp sequencing across 3 runs per sample was performed using the Illumina NextSeq 1000/2000 P2 reagents (200 cycles) v3 kit (Illumina). The RNA sequencing workflow is shown in **Figure 1**.

**Figure 1.**
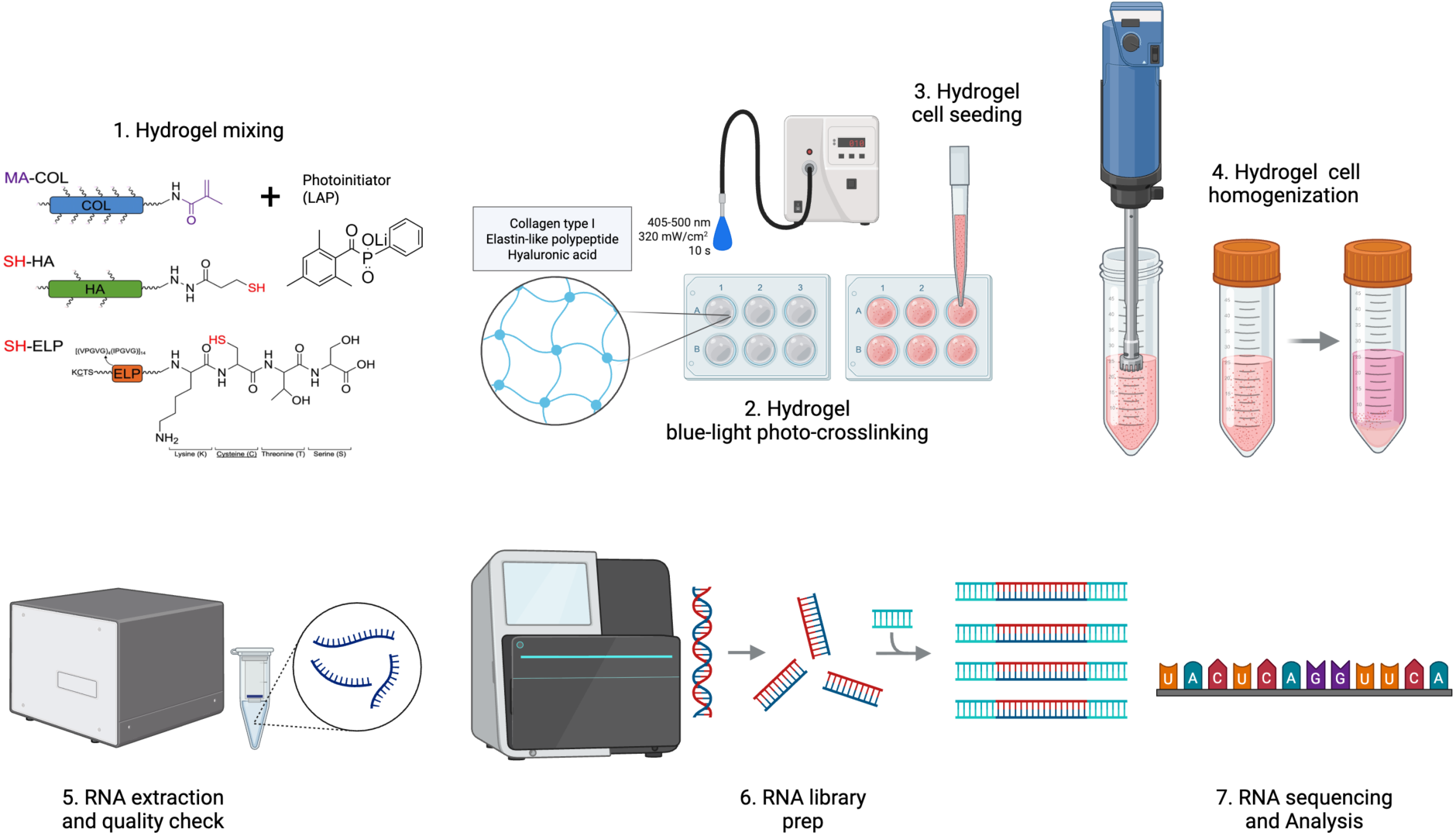
Biomimetic ECM hydrogel culture and RNA-sequencing workflow. (1) Methacrylate-conjugated collagen type I (MA-COL), thiol-conjugated hyaluronic acid (SH-HA), and thiol-containing elastin-like polypeptide (SH-ELP) were mixed with photoinitiator, lithium phenyl-2,4,6-trimethylbenzoylphosphinate (LAP); (2) dispensed into 6-well plates and blue-light photo-crosslinked (405-500 nm, 320 mW/cm², 10 s) into soft ECM hydrogels; (3) NTM and GTM cells were seeded on top; (4) cell-laden hydrogels were collected and homogenized in Trizol; (5) RNA was extracted and quality-checked; (6) libraries were prepared; and (7) sequenced (Illumina) and analyzed for differential expression, GO enrichment, and pathway networks. Created with BioRender.com

### 2.5 RNA sequencing analysis

RNA-seq data for cultured NTM and GTM cells (N = 5 each) were assessed for quality and analyzed using established pipelines on the Illumina Partek Flow platform (v.11.0.23.1105; Illumina), as previously described^40^. Briefly, sequence quality and coverage depth were assessed, reads were aligned to the human genome assembly (hg38) using STAR (v.2.7.8a), and aligned reads were annotated and quantified against the hg38 GENCODE Genes release 38 annotation. Counts were normalized using the median-ratio method as implemented in DESeq2, and differential expression analysis was performed with DESeq2, including four cofactors in the model: glaucoma status, originating laboratory of the TM cell strain, donor sex, and donor age. Differentially expressed genes (DEGs) with p < 0.05 and |fold change| ≥ 2 were considered significant. qRT-PCR analysis of select DEGs is described in the **Supplementary Information**.

Gene Ontology (GO) enrichment of the DEGs was performed in Partek Flow using the Gene Set Enrichment tool. GO terms with p < 0.05 in the Biological Process, Cellular Component, and Molecular Function categories were considered enriched. To examine how these terms were related, the DEG list was further analyzed using DAVID Functional Annotation Clustering (Knowledgebase v2026_2), which assembles GO terms that share genes. Guided by these clusters, the enriched terms were manually curated into three disease-relevant programs: (i) TGF-β/Wnt-associated ECM remodeling and fibrosis, (ii) Inflammation, and (iii) Hypoxia/metabolic stress. For each program, GO terms were selected based on their enrichment score and their relevance to TM biology and glaucoma. Only GO terms with two or more overlapping genes were considered. Redundant terms and terms describing unrelated tissues (e.g., heart development) were excluded.

To identify enriched pathways, gene ontologies, and candidate upstream regulators, the complete DEG list and the up-regulated subsets in GTM and NTM cells were uploaded into Ingenuity Pathway Analysis (IPA; Qiagen, Hilden, Germany) for Core Analysis to identify enriched networks/pathways, upstream regulators, and potential causal relationships. The RNA-seq dataset is being prepared for deposition in the NCBI Gene Expression Omnibus (GEO).

## Results

### 3.1 RNA-seq reveals distinct transcriptomic signatures in GTM cells compared to NTM cells in a physiologically-relevant ECM microenvironment

To examine glaucoma-associated transcriptional differences in a tissue-like soft ECM environment, we performed bulk RNA-seq on primary human NTM and GTM cells (N = 5 per group; mean NTM cell donor age: 66.8 years; mean GTM cell donor age: 68.4 years; **Table 1**) cultured on biomimetic ECM hydrogels^41,52^ that exhibited an elastic modulus of 0.42 ± 0.12 kPa (**Supplementary Figure 2**). This puts the bioengineered hydrogel stiffness close to the lower range of native TM tissue (0.9-16 kPa^25,36,37^), while simulating a more favorable physiologically-relevant microenvironment compared to supra-physiologically stiff conventional tissue culture plates.

Principal component analysis (PCA) showed significant variations among individual TM cell strains without clear separation by glaucoma status (**Figure 2A**). PC1 and PC2 explained 24.25% and 19.40% of the total variance, respectively. The relatively low percentage of total variance (43.64% combined) revealed the high heterogeneity and subtle biological difference of different strains of primary human GTM and NTM cells, which could be related to the ECM hydrogel culture environment. The lack of single dominant principal component suggests the lack of single dominant biological factor to explain the individual variations. Therefore, glaucoma status, originating laboratory, donor sex, and donor age were included as co-factors in the DESeq2 model for subsequent differential expression analysis.

**Figure 2.**
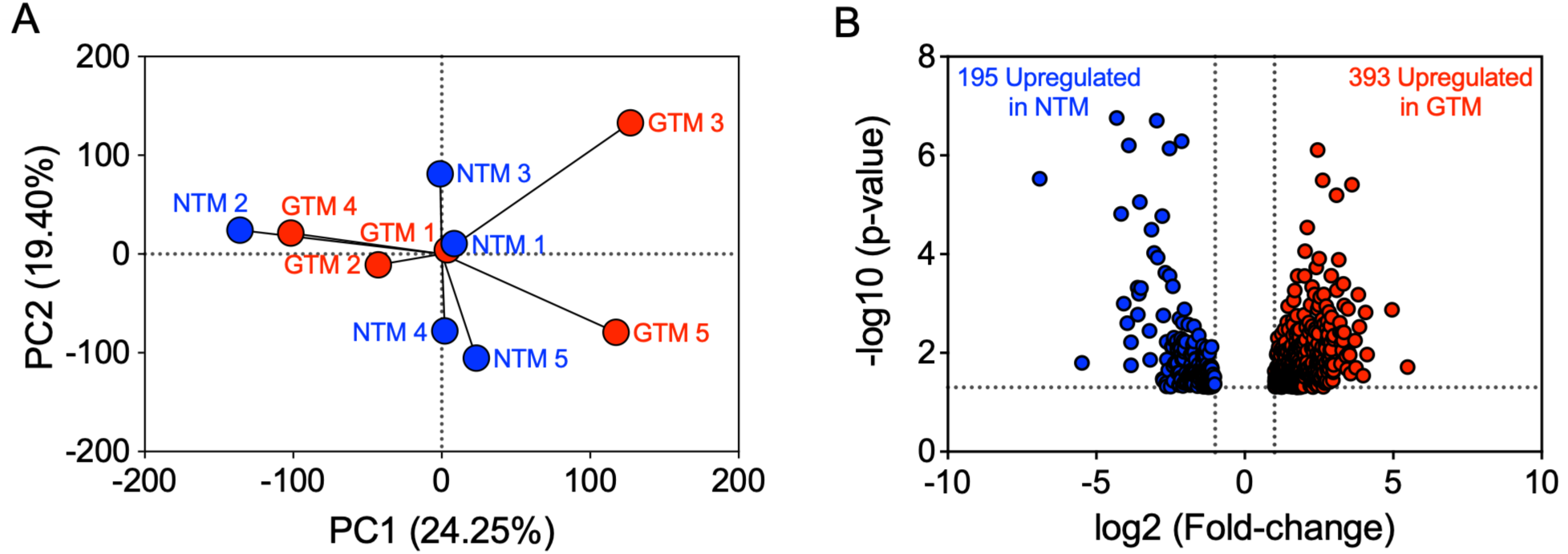
Bulk RNA-seq identifies distinct transcriptomic signatures in GTM cells compared to NTM cells cultured on biomimetic ECM hydrogels. (**A**) Principal component analysis of the RNA-seq samples colored by disease status. Each point represents one donor-derived cell strain (N = 5 per group); samples show variation without clear separation by glaucoma status. PC1 and PC2 account for 24.25% and 19.40% of the variance, respectively. (**B**) Volcano plot of the 588 DEGs in the GTM versus NTM comparison (p < 0.05; |fold change| ≥ 2), comprising 393 genes upregulated in GTM cells (red) and 195 genes downregulated in GTM cells (i.e., upregulated in NTM cells; blue).

Differential expression analysis identified 588 DEGs, comprising 393 genes upregulated in GTM cells and 195 genes downregulated in GTM cells (i.e., upregulated in NTM cells) (p < 0.05; |fold change| ≥ 2; **Figure 2B**).

We next investigated the top 15 differentially expressed genes in GTM and NTM cells (**Table 2**). The complete gene-level differential expression results are provided in **Supplementary Dataset 1**. The top genes upregulated in GTM cells encompass diverse functions in immune surveillance, metabolism, cell-matrix interactions, and cellular signaling. They are spanning several broad aspects of cellular homeostasis rather than defining certain functional pathways. In contrast, the top genes downregulated in GTM cells (i.e., upregulated in NTM cells) prominently represent ECM organization, tissue mechanics, and regulation of cellular adaptation.

**Table 2.**
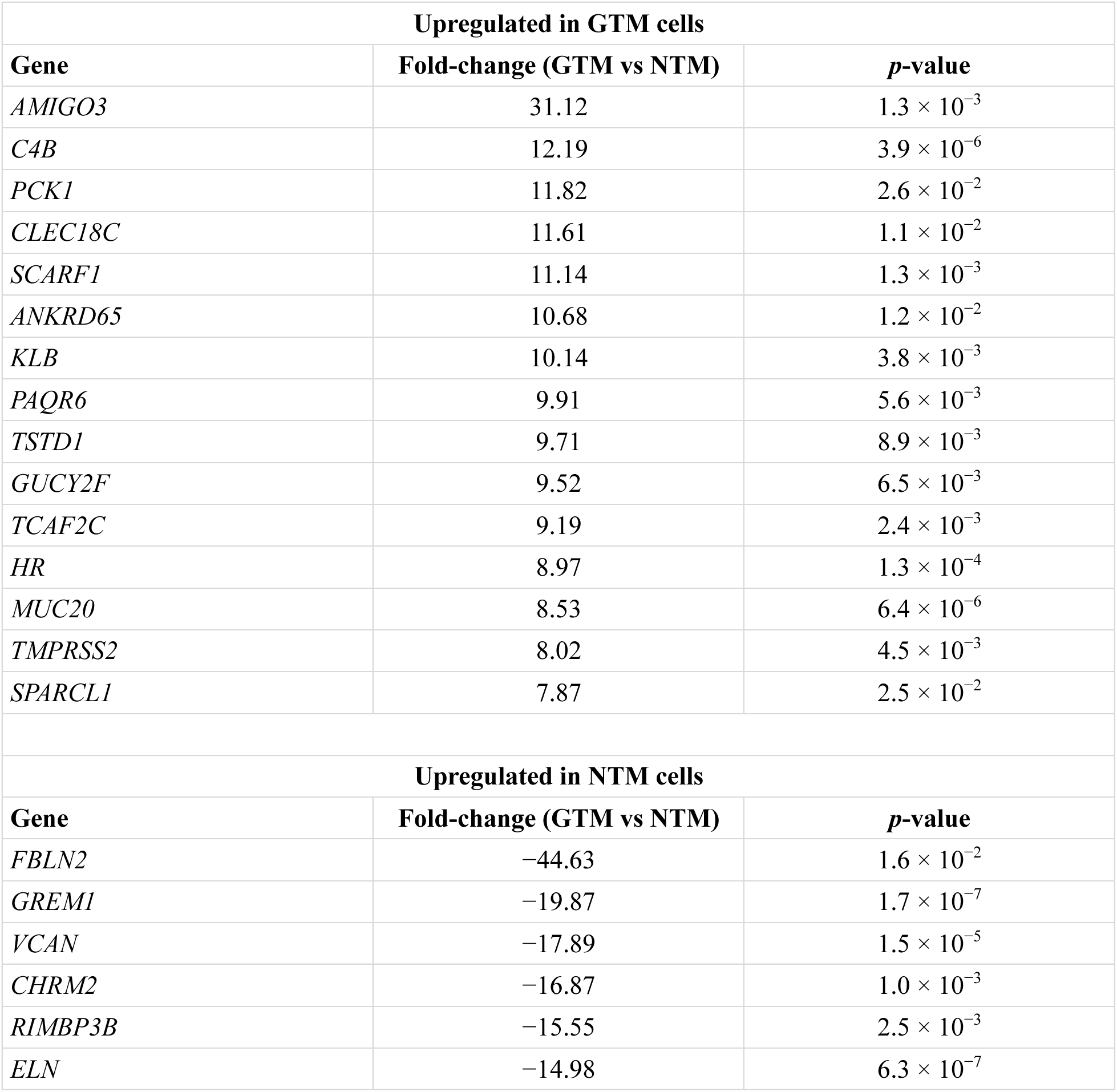
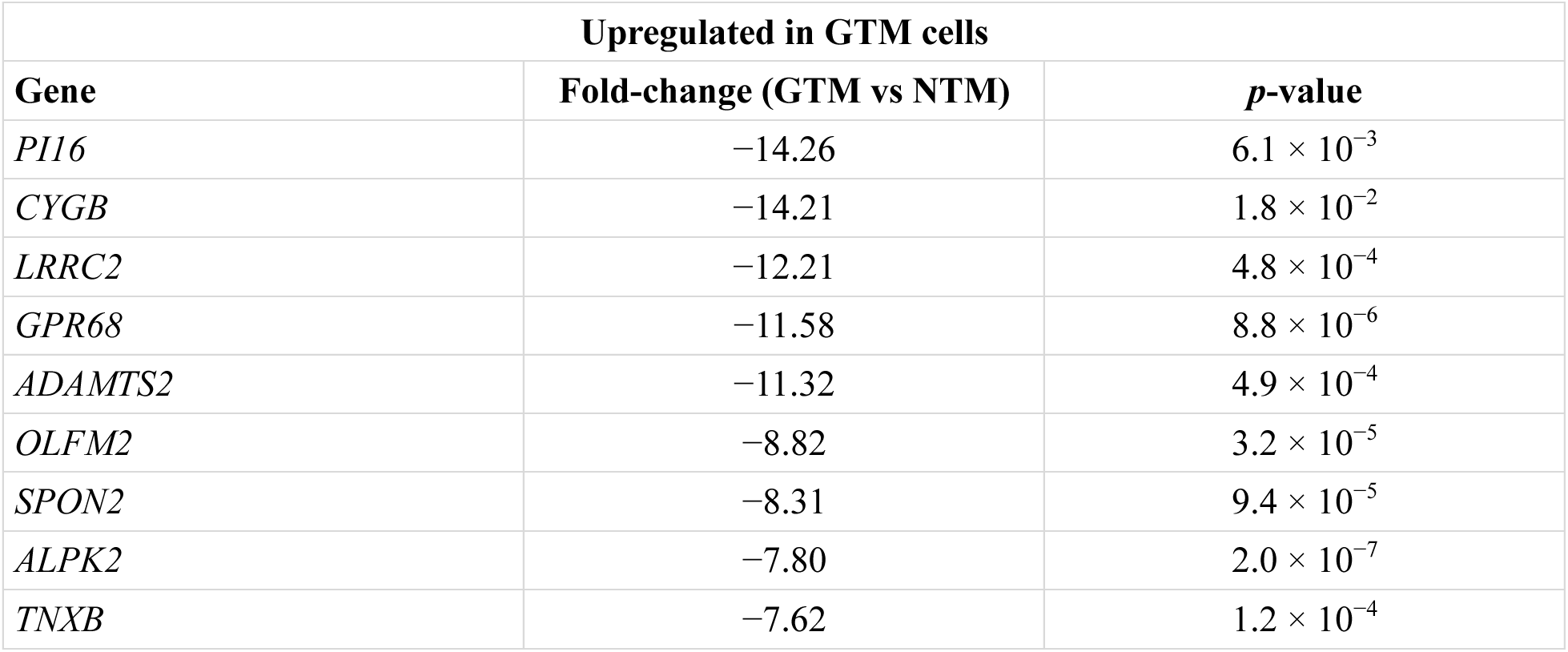
Top 15 differentially expressed genes in GTM and NTM cells cultured on biomimetic ECM hydrogels. Genes shown are the annotated protein-coding genes with the largest fold-change in each direction among DEGs meeting p < 0.05 and |fold change| ≥ 2. Fold-change is reported as GTM relative to NTM; negative values indicate higher expression in NTM cells.

Taken together, these data indicate that NTM and GTM cells maintain distinct ECM-remodeling-associated transcriptional profiles under common soft, ECM culture conditions. Notably, several of the ECM and fibrosis-associated genes that separate the two groups - including *ELN*, *VCAN*, *TNXB*, *ADAMTS2*, *COL1A1*, and *GREM1*-were expressed at lower levels in GTM cells than in NTM cells. These findings therefore support persistent, glaucoma-associated differences in ECM regulation rather than a uniform profibrotic transcriptional state in GTM cells.

Of note, qRT-PCR analysis of 10 DEGs - selected based on the magnitude and statistical significance of their expression changes as well as their functional relevance to ECM regulation and other glaucoma-associated cellular processes - reproduced the direction of change observed by RNA-seq but did not reach statistical significance (**Supplementary Figure 3**), with inherent donor variability and limited sample size likely driving this result.

### 3.2 Gene ontology enrichment organizes the GTM transcriptome into three disease programs: TGF-β/ECM remodeling, inflammation, and hypoxia/metabolism

To organize the 588 DEGs into biological programs, we performed gene ontology (GO) enrichment across the Biological Process, Cellular Component, and Molecular Function domains. Enriched terms were categorized into three disease-relevant programs: TGF-β/Wnt-associated ECM remodeling and fibrosis (**Table 3**), inflammation (**Table 4**), and hypoxia/metabolic stress (**Table 5**). The complete list of enriched terms and their program assignments is provided in **Supplementary Dataset 2**.

**Table 3.**
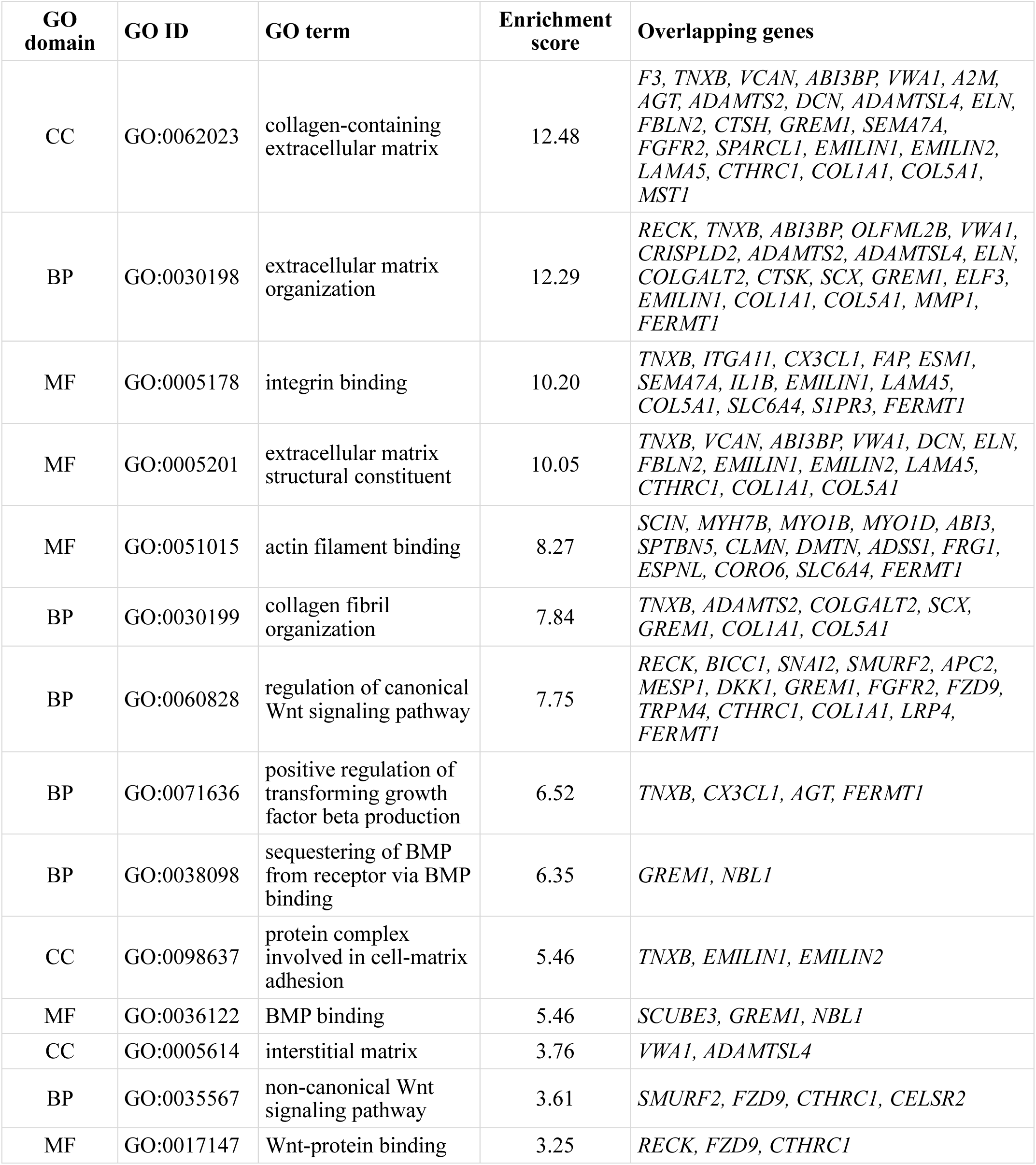
Gene Ontology enrichment terms associated with the disease program “TGF-β/Wnt-associated ECM remodeling and fibrosis”. GO terms with p < 0.05 in the categories Biological Process (BP), Cellular Component (CC), and Molecular Function (MF) were considered enriched and further analyzed for shared genes across GO terms using DAVID Functional Annotation Clustering.

**Table 4.** Gene Ontology enrichment terms associated with the disease program “Inflammation”. GO terms with p < 0.05 in the categories Biological Process (BP), Cellular Component (CC), and Molecular Function (MF) were considered enriched and further analyzed for shared genes across GO terms using DAVID Functional Annotation Clustering.

| GO domain | GO ID | GO term | Enrichment score | Overlapping genes |
| --- | --- | --- | --- | --- |
| MF | GO:0019955 | cytokine binding | 6.74 | *SCUBE3*, *CCR10*, *A2M*, *KIT*, *ACKR4*, *GREM1*, *LRRC32*, *NBL1*, *CRLF1* |
| BP | GO:0002720 | positive regulation of cytokine production involved in immune response | 6.66 | *SPON2*, *KIT*, *SEMA7A*, *HLA-A*, *IL1B*, *LACC1*, *ARID5A* |
| BP | GO:0006952 | defense response | 6.13 | *PRDM1*, *MACIR*, *STAT5A*, *NR4A1*, *SPON2*, *ABI3*, *ISG20*, *A2M*, *CX3CL1*, *C4A*, *C4B*, *BPGM*, *BRINP1*, *KIT*, *LAT*, *GPR68*, *KCNJ8*, *P2RY11*, *ELF3*, *GBP3*, *SEMA7A*, *HLA-A*, *HLA-C*, *IFI6*, *NDUFAF4*, *IL1B*, *NEDD4*, *IRF7*, *LACC1*, *ARID5A*, *BATF2*, *UNC13D*, *IFITM1*, *S1PR3*, *NOX5*, *DLEC1* |
| BP | GO:0002250 | adaptive immune response | 4.55 | *PRDM1*, *SCART1*, *LAT*, *KCNJ8*, *CTSH*, *HLA-A*, *HLA-C*, *IRF7*, *TRPM4*, *UNC13D* |
| MF | GO:0019956 | chemokine binding | 4.43 | *CCR10*, *A2M*, *ACKR4* |
| BP | GO:0006954 | inflammatory response | 4.21 | *MACIR*, *NR4A1*, *A2M*, *CX3CL1*, *C4A*, *C4B*, *BPGM*, *KIT*, *LAT*, *GPR68*, *KCNJ8*, *ELF3*, *SEMA7A*, *IL1B*, *LACC1*, *S1PR3* |
| CC | GO:0042612 | MHC class I protein complex | 3.76 | *HLA-A*, *HLA-C* |
| MF | GO:0140375 | immune receptor activity | 3.28 | *F3*, *CCR10*, *ACKR4*, *CTSH*, *CRLF1* |
| BP | GO:0030449 | regulation of complement activation | 3.26 | *A2M*, *IL1B* |
| MF | GO:0001848 | complement binding | 3.12 | *C4A*, *C4B* |

**Table 5.** Gene Ontology enrichment terms associated with the disease program “Hypoxia and metabolism”. GO terms with p < 0.05 in the categories Biological Process (BP), Cellular Component (CC), and Molecular Function (MF) were considered enriched and further analyzed for shared genes across GO terms using DAVID Functional Annotation Clustering.

| GO domain | GO ID | GO term | Enrichment score | Overlapping genes |
| --- | --- | --- | --- | --- |
| BP | GO:0015671 | oxygen transport | 10.71 | *BPGM*, *HBB*, *CYGB*, *HBG1* |
| MF | GO:0005344 | oxygen carrier activity | 7.68 | *HBB*, *CYGB*, *HBG1* |
| MF | GO:0034185 | apolipoprotein binding | 5.93 | *PCSK9*, *LCAT*, *LRP4* |
| BP | GO:0051902 | negative regulation of mitochondrial depolarization | 5.18 | *FZD9*, *IFI6* |
| BP | GO:0033993 | response to lipid | 4.13 | *BGLAP*, *RARA*, *STC2*, *NR4A1*, *GPBAR1*, *SPON2*, *A2M*, *CA9*, *BRINP1*, *ADCY6*, *DKK1*, *KCNJ8*, *CTSH*, *GBP3*, *FGFR2*, *GNG2*, *FIBIN*, *IL1B*, *ARID5A*, *COL1A1*, *SCNN1D*, *SLC6A4*, *PCK1*, *PCNA* |
| BP | GO:0042632 | cholesterol homeostasis | 3.65 | *TMEM97*, *PCSK9*, *EPHX2*, *LCAT*, *LIPG* |
| CC | GO:0043020 | NADPH oxidase complex | 3.12 | *NOXA1*, *NOX5* |

One dominant disease program emerging from the GO enrichment analysis was “TGF-β/Wnt/ECM fibrosis” (**Table 3**). Enriched terms included ECM organization, collagen fibril organization, collagen-containing ECM, integrin binding, and ECM structural constituents, together with canonical and non-canonical Wnt signaling, positive regulation of TGF-β production, and BMP binding and sequestration. Representative genes spanned matrix structural and remodeling components (*TNXB*, *VCAN*, *ABI3BP*, *VWA1*, *ADAMTS2*, *COL1A1*, *DCN*) and TGF-β/Wnt regulators (*RECK*, *SMURF2*, *SCX*, *GREM1*, *FZD9*, *CTHRC1*).

Another disease program involved “Inflammation” (**Table 4**), with enrichment of positive regulation of cytokine production, defense and inflammatory responses, adaptive immune response, complement activation and binding, MHC class I, and cytokine/chemokine/immune-receptor binding. Representative genes included *SPON2*, *IL1B*, *A2M*, *CX3CL1*, *NR4A1*, *CCR10*, *ACKR4*, *HLA-A*/*HLA-C*, and *C4A*/*C4B*.

A third prominent disease program involved “Hypoxia-and metabolism-related pathways” (**Table 5**). Enriched terms encompassed oxygen transport and carrier activity, negative regulation of mitochondrial depolarization, response to lipid, cholesterol homeostasis, NADPH oxidase complex, and apolipoprotein binding. Representative genes included the oxygen-handling genes *HBB*, *CYGB*, and *HBG1*, the stress-responsive nuclear receptor *NR4A1*, and the metabolic regulators *PCSK9*, *LCAT*, *NOX5*, and *PCK1*.

Together, these findings highlight three major disease-relevant themes within the glaucomatous TM transcriptome: ECM remodeling/fibrosis, inflammation and hypoxia/metabolic dysregulation.

### 3.3 Ingenuity pathway analysis nominates TGF-β, NF-κB, and p300/CBP as candidate upstream regulators of the three identified disease programs

Pathway-level analysis can organize differential gene expression into biologically relevant regulatory networks, including potential upstream regulators^53^. TGF-β signaling^22,54^, NF-κB-associated inflammatory activity^14,15,55,56^, and hypoxia-associated stress^16–19,57^ have each been proposed to contribute to TM dysfunction. However, these programs have generally been studied separately, and an integrated network view that links them in GTM cells profiled on biomimetic ECM hydrogels has not been described.

To identify candidate upstream regulators that organize these programs, the GTM DEGs were analyzed by Ingenuity Pathway Analysis (IPA) to maps causal networks within the framework of the three identified disease programs. The first program identified was a TGF-β-centered network linked to ECM remodeling (**Figure 3A**), with predicted inhibition of TGF-β based on the direction of its downstream targets including *FBLN2, GREM1, COL1A1, MMP1,* and *DCN*. The focused TGF-β family complex sub-network is shown in **Supplementary Figure 4A**.

**Figure 3.**
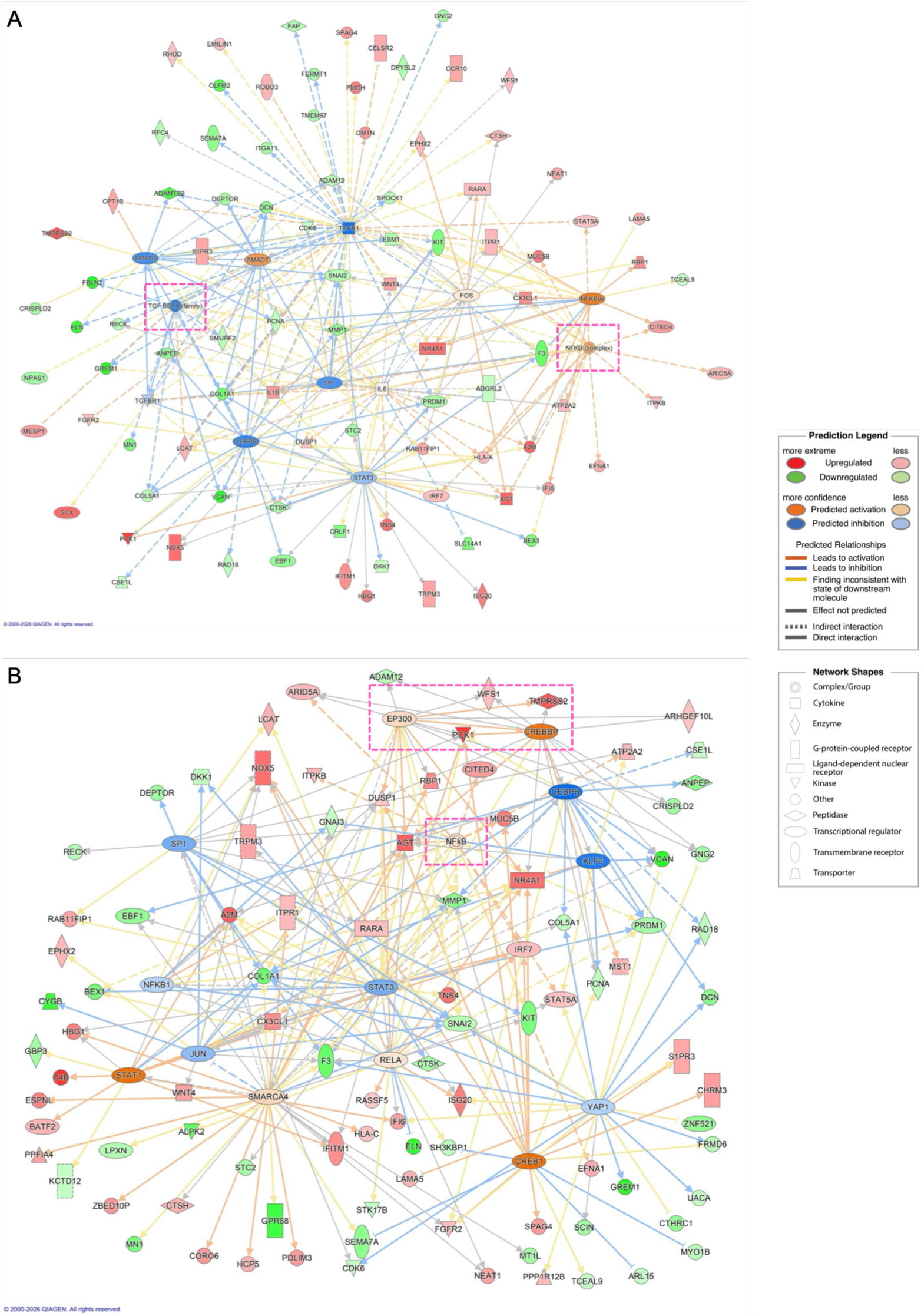
Ingenuity Pathway Analysis (IPA) regulatory networks linking TGF-β signaling to inflammatory/ECM targets and p300/CREB-associated transcriptional regulation in GTM cells compared to NTM cells cultured on biomimetic ECM hydrogels. IPA causal networks centered on (**A**) TGF-β and NF-κB signaling and (**B**) EP300 and CREB-family transcriptional regulators together with NF-κB inflammatory nodes, and their predicted downstream targets. The pink boxes highlight the three disease program nodes.

An NF-κB-associated inflammatory network was also identified, connecting NF-κB to inflammatory network components including *IL6, TNFSF10, CX3CL1*, and complement and HLA genes (**Figure 3A,B**). The focused NF-κB complex sub-network is shown in **Supplementary Figure 4B**.

The third program converged on a chromatin co-activator network centered on the histone acetyltransferases p300 (encoded by *EP300*) and CBP (encoded by *CREBBP*), together with *BRD4*, *DNMT1*, *KAT2B*, and *SMARCA4* and linked to ECM (*COL1A1*, *COL1A2*, *COL3A1*, *LOXL1*, *SERPINH1*) and inflammatory (IL6, NF-κB) targets (**Figure 3B**). In addition, hypoxia-associated stress responses and p300 co-activator activity emerged as a linked regulatory program via *NR4A1*, *PCK1*, and *DUSP1* (**Figure 3C**). The focused EP300 sub-network is shown in **Supplementary Figure 4C**.

## Discussion

ECM remodeling^7–13^, inflammation^14,15^, cellular stress^16–19^, and altered mechanotransduction^12,27,58–63^ have been associated with TM dysfunction in POAG, but the regulatory programs that sustain these phenotypes remain incompletely defined. Previous studies of the TM transcriptome, including their experimental systems, profiling methods, and principal findings, are summarized in **Supplementary Table 1.** The transcriptomic profiles discussed here identify three disease programs in GTM cells cultured on soft biomimetic hydrogels (elastic modulus, 0.42 ± 0.12 kPa): (i) TGF-β/Wnt-associated ECM remodeling, (ii) NF-κB-associated inflammation, and (iii) hypoxia/metabolic stress, with pathway analysis nominating p300/CBP chromatin co-activation as a candidate regulatory mechanism.

The first program is “TGF-β/Wnt-associated ECM remodeling”. Aberrant ECM deposition, crosslinking, and stiffening of the TM are central to elevated outflow resistance in POAG^7–13^. Our data show that glaucoma-associated transcriptional differences in ECM regulation remain detectable even after GTM cells are removed from their diseased tissue and cultured in the same soft biomimetic ECM environment as NTM cells. Thus, these differences do not require continued exposure to the native glaucomatous matrix to manifest during transcriptomic profiling – a key characteristic of TM cellular memory^64^. Cellular mechanical memory provides a framework for interpreting the persistence of disease-associated cellular signatures that persist even after the original pathological cue is removed. Cellular plasticity becomes altered through induction of stable alterations in chromatin organization, epigenetic modifications, and transcriptional programs that serve as nuclear repositories of prior experiences^65^. Our findings suggest that GTM cells may retain a transcriptional state shaped by their prior mechanical environment, even when cultured in a soft ECM environment. The nominated p300/CBP network further suggests a candidate connection between retained glaucomatous cellular identity and chromatin-dependent transcription.

Among the differentially expressed ECM genes, *ELN*, *VCAN*, *TNXB*, and the matrix-remodeling enzyme *ADAMTS2* contribute to matrix structure and turnover. Additional matricellular factors including *DCN*, *FBLN2*, and *SPON2* further implicate altered matrix regulation^9–11^. *DCN* provides a functional link between matrix regulation and TGF-β-associated outflow dysfunction. Schneider et al. found that *DCN* deficiency increased IOP and that soluble *DCN* attenuated TGF-β2-induced ECM-related responses in human TM cells^66^. Hill et al. demonstrated that recombinant decorin reduced established TM fibrosis and lowered IOP in a TGF-β-induced rodent model^67^. The lower expression of *ELN*, *ADAMTS2*, *COL1A1*, and *GREM1* in GTM cells suggests that the ECM-associated signature reflects altered matrix composition and regulation rather than a coordinated increase in all fibrosis-related transcripts. *SCX*, which encodes a transcription factor that regulates collagen-associated fibrotic programs^68,69^, and *FAP*, which encodes a marker of activated fibroblasts in remodeling tissues^70,71^, were also altered. TGF-β2 is elevated in the aqueous humor of glaucomatous eyes and promotes ECM accumulation, cytoskeletal reorganization, and contractile remodeling in TM cells^13,17,20–22,38^. Several DEGs connect directly to this axis. *SMURF2*, which encodes a ubiquitin ligase that regulates SMAD turnover and the amplitude of TGF-β signaling^72–75^, and *VEPH1*, which modulates TGF-β signal strength^76,77^, are both differentially expressed. *WNT4* further connects this network to the Wnt pathway, and an imbalance between Wnt and TGF-β signaling has been associated with TM dysfunction and elevated IOP^78–83^. *GREM1* encodes a BMP antagonist that can shift the BMP/TGF-β balance toward a profibrotic state. *GREM1* was previously reported to be elevated in GTM cells and to promote TGF-β-associated ECM production^84,85^. Our data contextualizes these prior observations through the lens of genome-wide transcriptomics. Of note, a recent proteomic profiling study of primary NTM and GTM cells identified altered ECM, Wnt, cytoskeletal, protein-degradation, and nuclear pathways^86^. These protein-level findings support persistent molecular differences in cultured GTM cells and provide a complementary framework for interpreting disease-associated transcriptional changes.

The second identified program is “Inflammation”. Chronic, low-grade inflammation has been increasingly recognized in the anterior segment of glaucomatous eyes, including the TM^56,87–89^. Our transcriptomic data extend these observations by identifying inflammatory signatures among the DEGs in GTM cells. Enriched immune terms included cytokine-production and inflammatory-response genes (*IL1B*, *SPON2*, *A2M*, *CX3CL1*, and *NR4A1*), chemokine and complement components (*CCR10*, *ACKR4*, *C4A*, *C4B*), and MHC class I genes (*HLA-A*, *HLA-C*). Differentially expressed *ARID5A* encodes an RNA-binding protein that stabilizes inflammatory cytokine mRNAs and can amplify NF-κB-associated responses post-transcriptionally^89–94^.The long non-coding RNA *ZNF295-AS1* further suggests that non-coding regulatory layers may contribute to the inflammatory and post-transcriptional landscape of GTM cells^95–100^, although its role in TM remains to be established.

The third program involves “Hypoxia/metabolic stress”. Hypoxia-associated stress and metabolic dysfunction have been described in glaucoma and may contribute to the persistent stress phenotype of GTM cells^57,101–103^. Among the DEGs, we observed oxygen-transport and carrier genes (*HBB*, *CYGB*, *HBG1*), the stress-responsive regulator *NR4A1*^104–106^, and genes involved in lipid and glucose metabolism (*PCSK9*, *LCAT*, *PCK1*). Up-regulation of *ATP2A2*, which encodes the SERCA2 calcium pump^107^, implicates altered ER calcium homeostasis, consistent with reports of ER stress in GTM cells^108–112^. The long non-coding RNA *NEAT1*, a hypoxia-and inflammation-responsive paraspeckle scaffold, may provide a non-coding regulatory link to these stress signatures^103,113–116^. These changes are also consistent with reports of mitochondrial deficits in GTM cells^117–125^.

Epigenetic co-activation may help explain how GTM cells sustain the ECM, inflammatory, and stress programs identified above^126,127^. Pathway analysis nominated the histone acetyltransferases p300 (*EP300*) and CBP (*CREBBP*), with additional chromatin regulators, as a candidate co-activator network. In fibrotic disorders such as systemic sclerosis and cardiac fibrosis induced by TGF-β, p300 is recruited to ECM gene promoters including *COL1A2*, driving histone H3/H4 acetylation and collagen induction^128–132^. p300 also serves as a co-activator for HIF-1α and can integrate TGF-β, hypoxia, and inflammatory signaling at the chromatin level^130,133–136^. In TM cells, the role of p300 remains incompletely defined. Prior work has shown that global protein and histone H3 acetylation differs between NTM and GTM cells, and that histone hyperacetylation is sufficient to drive TGF-β2 and ECM protein expression in TM cells and to elevate IOP in perfused eyes^137^. These earlier studies position the histone acetylation balance as a candidate upstream driver of GTM cell dysfunction, but the specific acetyltransferases responsible are unknown. The p300/CBP network identified by our pathway analysis is consistent with these observations and nominates p300 as a candidate upstream regulator that could integrate the ECM/fibrosis, inflammatory, and hypoxia/metabolic programs^126,130,134^. The predicted p300/CBP network does not demonstrate altered p300 protein function. Direct measurement of p300 abundance, chromatin occupancy, and histone acetylation activity, together with functional perturbations, will be needed to test this regulatory model in GTM cells.

In summary, a central feature of our study is that it extends prior NTM and GTM transcriptomic comparisons by examining five independent primary donor cell strains per group from three laboratories on a common biomimetic ECM substrate. The mean ECM hydrogel elastic modulus of ∼0.4 kPa is positioned close to the lower end of native TM tissue, ranging from 0.9 to 16 kPa^25,36,37^, while simulating a more favorable tissue-like soft ECM microenvironment compared to stiff conventional tissue culture plates. Using this unique study design, we identified novel glaucoma-associated transcriptional programs that provide both a reference for primary human GTM cells and a starting point for functional validation of ECM, inflammatory, and stress-associated regulators in TM dysfunction.

## Limitations

Some limitations should be considered when interpreting these findings. The cohort is limited by a relatively small sample size (N = 5 cell strains each) and heterogeneity in age, sex, glaucoma subtype, and IOP-lowering medication exposure (see **Table 1**). POAG or glaucoma status categorization of the postmortem donor tissues was based on limited eye bank records that may increase the heterogeneity of the included TM cells. Principal component analysis showed variation among cell strains without clear separation by glaucoma status. Donor sex, donor age, originating laboratory, and glaucoma status were thus included in the DESeq2 model, although residual effects of donor and culture differences cannot be excluded. qRT-PCR reproduced the principal direction of change for the tested genes, but none of the comparisons reached statistical significance. The modest cohort and inter-strain variability likely played a role.

## Conclusion

Primary GTM cells retain glaucoma-associated ECM remodeling, inflammatory, and stress-response programs after removal from the diseased tissue and culture on soft ECM substrates – a hallmark of cellular mechanical memory^64^. These findings support investigating persistent cellular adaptations in GTM cell and tissue dysfunction and nominate p300/CBP as a candidate mechanism for functional testing.

## Future Directions

A central question arising from these data is whether modulating p300 can reverse glaucoma-associated changes in ECM regulation, inflammation, and hypoxia/metabolic signaling. Several approaches would directly test the programs identified here. Genetic and/or pharmacologic loss-of-function studies could test whether reducing p300 attenuates the disease-associated phenotype in GTM cells, while gain-of-function approaches would test whether increasing p300 in NTM cells induces pathologic changes similar to GTM cells to establish causality. Readouts would include histone acetylation, nuclear organization, ECM deposition, cell stiffness, contractility, and inflammatory and metabolic responses. Chromatin immunoprecipitation sequencing or CUT&RUN epigenomic mapping of GTM and NTM cells on biomimetic hydrogels would assess p300 occupancy and histone acetylation to test the chromatin mechanism. Integrated single-cell/nucleus ATAC-seq and RNA-seq analysis would link transcriptional programs to the underlying chromatin regulatory landscape, and resolve whether the disease programs are uniform across the TM population or concentrated in particular subtypes^138^. Larger donor cohorts will help separate persistent disease features from inter-individual variability.

## Disclosure

The authors report no conflicts of interest.

## Funding

This research was supported by grants from the NIH (R01EY034096 (SH), R21EY036189 (SH), R01EY032960 (YL), R21EY033961 (YL), R01EY019643 (KEK), R01EY032590 (KEK), R01EY028608 (WDS), R01EY031710 (WDS), P30EY005722 (WDS), R01EY036880 (PSG), the BrightFocus Foundation (G2024006S), the Glaucoma Research Foundation (YL), The Glaucoma Foundation (YL), Research to Prevent Blindness (Career Advancement Award (SH) and unrestricted grant to SUNY Upstate Medical University Department of Ophthalmology and Visual Sciences), and Lions Region 20-Y1 (unrestricted grant to SUNY Upstate Medical University Department of Ophthalmology and Visual Sciences).

## Acknowledgments

We thank Drs. Robert W. Weisenthal, Matthew Weinstein, and Stephen G. Spitzer and the team at Specialty Surgery Center of Central New York for assistance with corneal rim specimens, and VisionGift, Portland, OR for help procuring human donor eyes. We also thank the SUNY Upstate Medical University Molecular Analysis Core for helping in RNA library preparation and sequencing.

## Author contributions

R.G. and S.H. designed all experiments and collected the data. R.H. and Y.L. analyzed the data. A.N.S. helped in analyzing the data. K.E.K. and W.D.S. provided certain NTM and GTM cells. R.G. and S.H. wrote the manuscript. All authors commented on and approved the final manuscript. P.S.G. and S.H. conceived and supervised the research.

## Data and materials availability

All data needed to evaluate the conclusions in the paper are present in the paper and/or the Supplementary Information. Additional data related to this paper may be requested from the authors.

## Notes

### Competing Interest Statement

The authors have declared no competing interest.

