## Supplementary Information for "Transcriptomic profiling of normal and glaucomatous trabecular meshwork cells in a tissue-like soft ECM environment"

***TM cell characterization***

HTM 28 cells were seeded at $2 \times 10^4$ cells/cm$^2$ in 6-well culture plates or on sterilized glass coverslips in 24-well culture plates, and cultured in DMEM with 10% FBS and 1% PSG[1]. TM cell morphology and growth characteristics were monitored by phase contrast microscopy using an LMI-3000 Series Routine Inverted Microscope (Laxco; Thermo Fisher Scientific). Once confluent, TM cells were treated with 100 nM dexamethasone (DEX; Fisher Scientific) or vehicle control (0.1% (v/v) ethanol) in DMEM with 1% FBS and 1% PSG for 4 d, and serum- and phenol red-free DMEM for 3 d. The TM cell culture supernatants were collected and concentrated using Amicon® Ultra Centrifugal Filters (Millipore Sigma, Burlington, MA, USA) for immunoblot analysis. The monolayer TM cells were processed for immunocytochemistry analyses. DEX-induced myocilin (MYOC) upregulation in more than 50% of HTM cells was used as inclusion/exclusion criterion. Immunoblot analysis: Equal volumes of concentrated TM cell culture supernatants ± DEX at 7 d, supplemented with Halt™ protease/phosphatase inhibitor cocktail (Thermo Fisher Scientific), were mixed with 4× loading buffer (Invitrogen; Thermo Fisher Scientific) and 5% beta-mercaptoethanol (Fisher Scientific), boiled for 5 min, subjected to SDS-PAGE using NuPAGE™ 4-12% Bis-Tris Gels (Invitrogen; Thermo Fisher Scientific) at 150V for 120 min, and transferred to 0.45 µm PVDF membranes (Millipore Sigma). Membranes were blocked with 5% bovine serum albumin (Thermo Fisher Scientific) in tris-buffered saline with 0.2% Tween®20 (Thermo Fisher Scientific), and probed with a primary antibody against MYOC (mouse anti-MYOC; MABN866; 1:2,000; Sigma-Aldrich; St. Louis, MO, USA) followed by incubation with a fluorescent secondary antibody (IRDye® 800CW goat anti-mouse IgG; 926-32210; 1:15,000; LI-COR, Lincoln, NE, USA). Bound antibodies were visualized with an Odyssey CLx imager (LI-COR). Immunocytochemistry analysis: TM cells ± DEX at 7 d were fixed with 4% paraformaldehyde (Thermo Fisher Scientific) at room temperature for 10 min, permeabilized with 0.5% Triton™ X-100 (Thermo Fisher Scientific), blocked with normal goat serum (Abcam; Waltham, MA, USA), and incubated with a primary antibody against MYOC (rabbit anti-MYOC; ab41552; 1:200; Abcam) followed by incubation with a fluorescent secondary antibody (Alexa Fluor®488 goat anti-rabbit IgG; ab150077; Abcam). Nuclei were counterstained with 4',6'-diamidino-2-phenylindole (DAPI; Abcam). Coverslips were mounted with ProLong™ Gold Antifade (Thermo Fisher Scientific) on Superfrost™ Plus microscope slides (Fisher Scientific), and fluorescent images were acquired with an Eclipse Ni microscope (Nikon). Four fields of view per sample were analyzed to quantify percent of cells expressing MYOC, followed by calculation of mean +/- SD as follows:

DEX-induced MYOC upregulation = $(\text{Mean}_{DEX} - \text{Mean}_{Ctrl}) \pm [(\text{SD}_{DEX})2 + (\text{SD}_{Ctrl})^2]^{1/2}$.

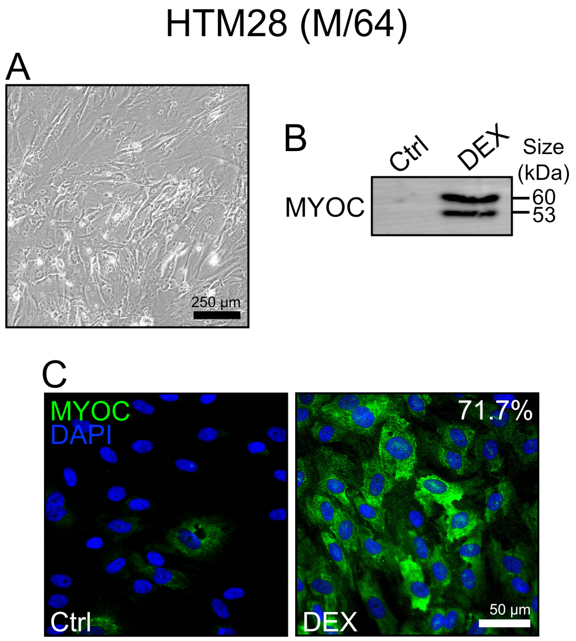


**Supplementary Figure 1. HTM28 cell characterization.** (**A**) Representative phase contrast micrograph. (**B**) Immunoblot of secreted myocilin (MYOC) in control (Ctrl) and dexamethasone-treated (DEX) cells at 7 d. (**C**) Representative fluorescence micrographs of intracellular MYOC in control and DEX-treated cells at 7 d (MYOC = green; DAPI = blue).

***ECM hydrogel rheology analysis***

Two hundred and fifty microliters of acellular hydrogel precursor solutions were pipetted into custom 16x1-mm PDMS molds[1] and crosslinked by exposure to blue light[2] (405–500 nm, 320 mW/cm$^2$, 10 s; OmniCure S1500 UV Spot Curing System; Excelitas Technologies, Mississauga, Ontario, Canada). A Kinexus rheometer (Malvern Panalytical, Westborough, MA, USA) fitted with an 8-mm diameter parallel plate was used to measure hydrogel viscoelasticity. To ensure standard conditions across all samples (N = 15), the geometry was lowered into the hydrogels until a calibration normal force of 0.02 N was achieved. Subsequently, an oscillatory shear-strain sweep test (0.1-60%, 1.0 Hz, 25°C) was applied to determine storage modulus (G') and loss modulus (G") in the linear region. Elastic modulus was calculated with $E = 2 * (1 + v) * G'$, where a Poisson's ratio (v) of 0.5 for the ECM hydrogels was assumed[3].

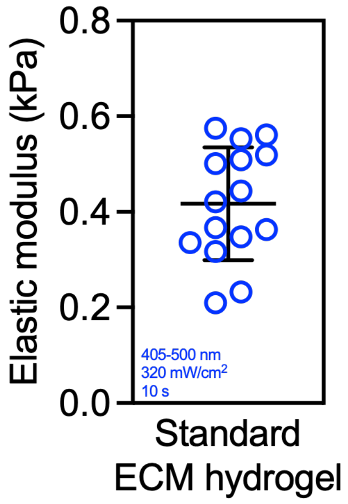


**Supplementary Figure 2. ECM hydrogel stiffness.** ECM hydrogel elastic modulus by oscillatory rheology analysis using standard ECM hydrogel formulation and photo-crosslinking conditions (N=15; Mean ± SD).

***Quantitative reverse transcription-polymerase chain reaction (qRT-PCR) analysis***

Total RNA was extracted from hydrogels containing newly cultured NTM/GTM cells maintained under the same conditions as those used for RNA-seq, and quality control was conducted by the Molecular Analysis Core. Three independent cell strains per group were used: HTM 36, HTM 28, and NTM 2022-0794; and GTM 0949, GTM 1672, and GTM 1898. GTM 211 and GTM 1445 were not included due to insufficient cell stocks. RNA concentration and integrity were assessed using an Agilent 2100 Bioanalyzer (**Supplementary Table 3**). One thousand ng of total RNA per sample were used for reverse transcription with iScript™ cDNA Synthesis Kit (BioRad, Hercules, CA, USA). One hundred ng of cDNA per reaction were amplified in technical triplicates for 40 cycles using a CFX 384 Real Time PCR System (BioRad), with an annealing temperature of 60°C, Power SYBR™ Green PCR Master Mix (Thermo Fisher Scientific), and custom-designed qRT-PCR primers for target genes (IDT, Coralville, IA, USA; **Supplementary Table 4**). Transcript levels were normalized to *GAPDH*, and GTM mRNA fold-changes were calculated relative to the average of NTM controls using the comparative CT method[4]. Differences in mRNA expression between NTM and GTM groups were assessed using an unpaired t-test in GraphPad Prism software v11.1 (GraphPad Software, La Jolla, CA, USA). Ten DEGs ($p < 0.05$; $|\text{fold change}| \geq 2$) were selected for qRT-PCR analyses, five up-regulated in GTM cells and five up-regulated in NTM cells (i.e., down-regulated in GTM cells), prioritized by the magnitude of the RNA-seq fold-change and the significance of the p-value, and restricted to genes with established relevance to ECM regulation or other glaucoma-associated cellular processes: *AMIGO3, C4B, CLMN, ELF3, ARID5A* (up-regulated in GTM cells) and *GREM1, ELN, ADAMTS2, KIT, F3* (up-regulated in NTM cells).

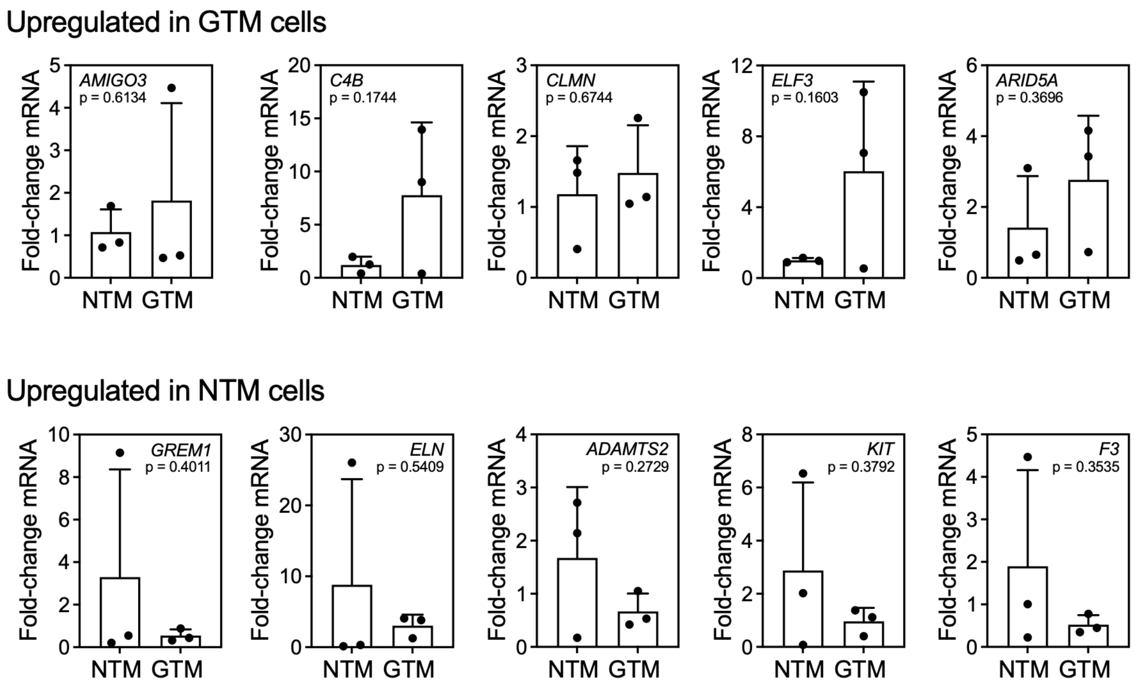


**Supplementary Figure 3. Select qRT-PCR gene expression analysis.** Normalized mRNA fold-changes by qRT-PCR (N=3 NTM/GTM cell strains; Mean ± SD).

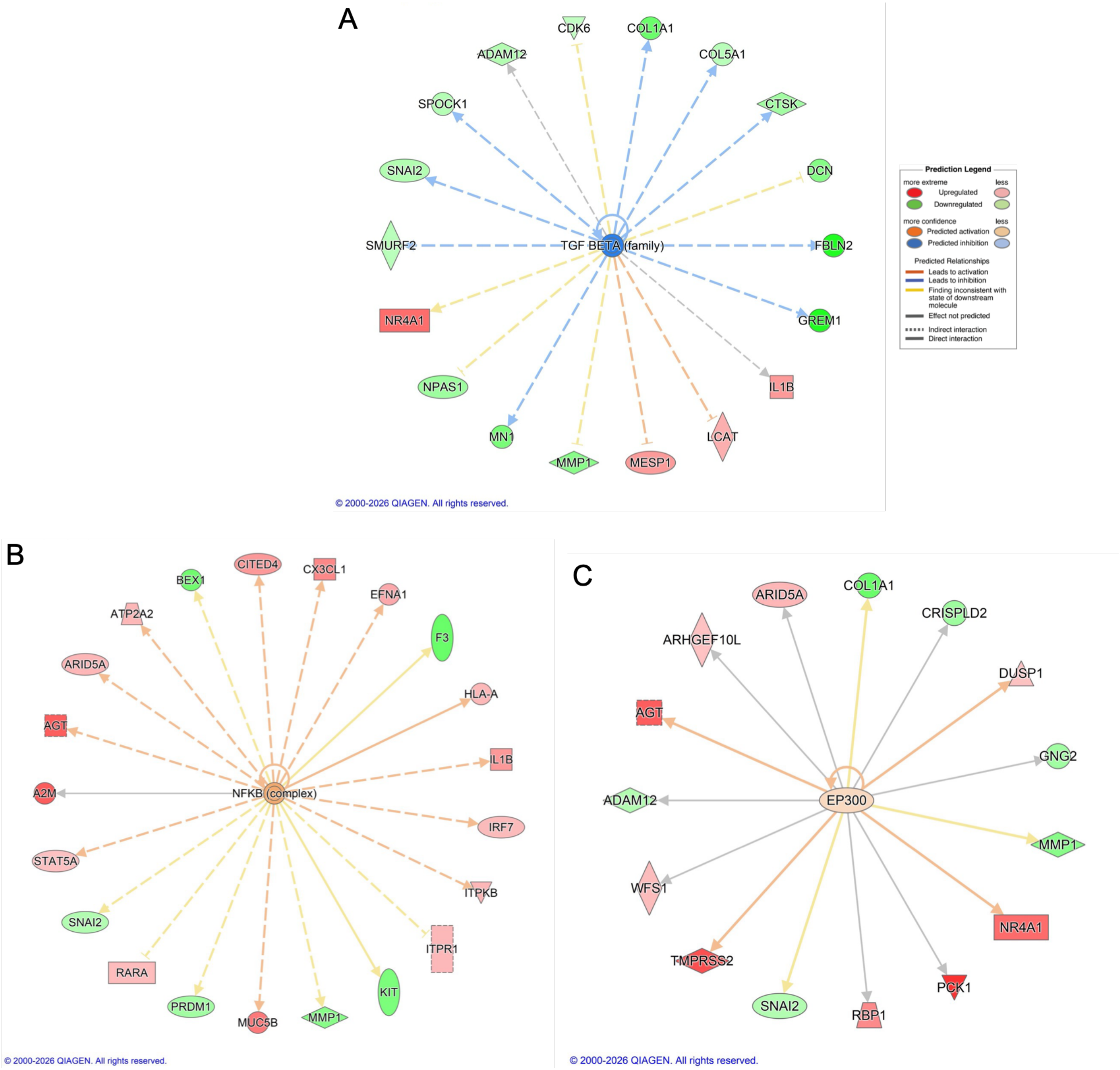


**Supplementary Figure 4. Focused Ingenuity Pathway Analysis (IPA) regulatory sub-networks in GTM cells compared to NTM cells cultured on biomimetic ECM hydrogels.** (**A**) **TGF-β family complex;** TGF-β is predicted to be inhibited, with downstream ECM- and fibrosis-related genes *COL1A1, COL5A1, DCN, FBLN2, GREM1, CTSK,* and *SNAI2* showing higher expression in NTM cells in green. (**B**) **NF-κB complex;** NF-κB is predicted to be activated, with downstream inflammation- and immune-related genes (*IL1B, CX3CL1, HLA-A, IRF7, STAT5A,* and *A2M*) showing higher expression in GTM cells in red. (**C**) **EP300 node;** EP300 is predicted to be activated, with downstream metabolic stress- and immunomodulatory-related genes (*AGT, NR4A1, TMPRSS2, PCK1,* and *DUSP1*) showing higher expression in GTM cells in red.
